# DeepMicro: deep representation learning for disease prediction based on microbiome data

**DOI:** 10.1101/785626

**Authors:** Min Oh, Liqing Zhang

**Affiliations:** Department of Computer Science, Virginia Tech, Blacksburg, VA, USA

## Abstract

Human microbiota plays a key role in human health and growing evidence supports the potential use of microbiome as a predictor of various diseases. However, the high-dimensionality of microbiome data, often in the order of hundreds of thousands, yet low sample sizes, poses great challenge for machine learning-based prediction algorithms. This imbalance induces the data to be highly sparse, preventing from learning a better prediction model. Also, there has been little work on deep learning applications to microbiome data with a rigorous evaluation scheme. To address these challenges, we propose DeepMicro, a deep representation learning framework allowing for an effective representation of microbiome profiles. DeepMicro successfully transforms high-dimensional microbiome data into a robust low-dimensional representation using various autoencoders and applies machine learning classification algorithms on the learned representation. In disease prediction, DeepMicro outperforms the current best approaches based on the strain-level marker profile in five different datasets. In addition, by significantly reducing the dimensionality of the marker profile, DeepMicro accelerates the model training and hyperparameter optimization procedure with 8X-30X speedup over the basic approach. DeepMicro is freely available at https://github.com/minoh0201/DeepMicro.

## Introduction

As our knowledge of microbiota grows, it becomes increasingly clear that the human microbiota plays a key role in human health and diseases ^1^. The microbial community, composed of trillions of microbes, is a complex and diverse ecosystem living on and inside a human. These commensal microorganisms benefit humans by allowing them to harvest inaccessible nutrients and maintain the integrity of mucosal barriers and homeostasis. Especially, the human microbiota contributes to the host immune system development, affecting multiple cellular processes such as metabolism and immune-related functions ^1,2^. They have been shown to be responsible for carcinogenesis of certain cancers and substantially affect therapeutic response ^3^. All these emerging evidences substantiate the potential use of microbiota as a predictor for various diseases ^4^.

The development of high-throughput sequencing technologies has enabled researchers to capture a comprehensive snapshot of the microbial community of interest. The most common components of the human microbiome can be profiled with 16S rRNA gene sequencing technology in a cost-effective way ^5^. Comparatively, shotgun metagenomic sequencing technology can provide a deeper resolution profile of the microbial community at the strain level ^6,7^. As the cost of shotgun metagenomic sequencing keeps decreasing and the resolution increasing, it is likely that a growing role of the microbiome in human health will be uncovered from the mounting metagenomic datasets.

Although novel technologies have dramatically increased our ability to characterize human microbiome and there is evidence suggesting the potential use of the human microbiome for predicting disease state, how to effectively utilize the human microbiome data faces several key challenges. Firstly, effective dimensionality reduction that preserves the intrinsic structure of the microbiome data is required to handle the high dimensional data with low sample sizes, especially the microbiome data with strain-level information that often contain hundreds of thousands of gene markers but for only some hundred or fewer samples. With a low number of samples, large number of features can cause the curse of dimensionality, usually inducing sparsity of the data in the feature space. Along with traditional dimensionality reduction algorithms, autoencoder that learns a low-dimensional representation by reconstructing the input ^8^ can be applied to exploit microbiome data. Secondly, given the fast amounting metagenomic data, there is an inadequate effort in adapting machine learning algorithms for predicting disease state based on microbiome data. In particular, deep learning is a class of machine learning algorithms that builds on large multi-layer neural networks, and that can potentially make effective use of metagenomic data. With the rapidly growing attention from both academia and industry, deep learning has produced unprecedented performance in various fields, including not only image and speech recognition, natural language processing, and language translation but also biological and healthcare research ^9^. A few studies have applied deep learning approaches to abundance profiles of the human gut microbiome for disease prediction ^10,11^. However, there has been no research utilizing strain-level profiles for the purpose. Comparatively, strain level profiles, often containing hundreds of thousands of gene markers’ information, should be more informative for accurately classifying the samples into patient and healthy control groups across different types of diseases than abundance profiles that usually contain only a few hundred bacteria’s abundance information ^12^. Lastly, to evaluate and compare the performance of machine learning models, it is necessary to introduce a rigorous validation framework to estimate their performance over unseen data. Pasolli et al., a study that built classification models based on microbiome data, utilized a 10-fold cross-validation scheme that tunes the hyper-parameters on the test set without using a validation set ^12^. This approach may overestimate model performance as it exposes the test set to the model in the training procedure ^13,14^.

To address these issues, we propose DeepMicro, a deep representation learning framework that deploys various autoencoders to learn robust low-dimensional representations from high-dimensional microbiome profiles and trains classification models based on the learned representation. We applied a thorough validation scheme that excludes the test set from hyper-parameter optimization to ensure fairness of model comparison. Our model surpasses the current best methods in terms of disease state prediction of inflammatory bowel disease, type 2 diabetes in the Chinese cohort as well as European women cohort, liver cirrhosis, and obesity. DeepMicro is open-sourced and publicly available software to benefit future research, allowing researchers to obtain a robust low-dimensional representation of microbiome profiles with user-defined deep architecture and hyper-parameters.

## Methods

### Dataset and Extracting Microbiome Profiles

We considered publicly available human gut metagenomic samples of six different disease cohorts: inflammatory bowel disease (IBD), type 2 diabetes in European women (EW-T2D), type 2 diabetes in Chinese (C-T2D) cohort, obesity (Obesity), liver cirrhosis (Cirrhosis), and colorectal cancer (Colorectal). All these samples were derived from whole-genome shotgun metagenomic studies that used Illumina paired-end sequencing technology. Each cohort consists of healthy control and patient samples as shown in Table 1. IBD cohort has 25 individuals with inflammatory bowel disease and 85 healthy controls ^15^. EW-T2D cohort has 53 European women with type 2 diabetes and 43 healthy European women ^16^. C-T2D cohort has 170 Chinese individuals with type 2 diabetes and 174 healthy Chinese controls ^17^. Obesity cohort has 164 obese patients and 89 non-obese controls ^18^. Cirrhosis cohort has 118 patients with liver cirrhosis and 114 healthy controls ^19^. Colorectal cohort has 48 colorectal cancer patients and 73 healthy controls ^20^. In total, 1,156 human gut metagenomic samples, obtained from MetAML repository ^12^, were used in our experiments.

**Table 1.** Human gut microbiome datasets used for disease state prediction

| Disease | Dataset name | # total samples | # of healthy controls | # of patient samples | Data source references |
| --- | --- | --- | --- | --- | --- |
| Inflammatory Bowel Disease | IBD | 110 | 85 | 25 | <sup>15</sup> |
| Type 2 Diabetes | EW-T2D | 96 | 43 | 53 | <sup>16</sup> |
|  | C-T2D | 344 | 174 | 170 | <sup>17</sup> |
| Obesity | Obesity | 253 | 89 | 164 | <sup>18</sup> |
| Liver Cirrhosis | Cirrhosis | 232 | 114 | 118 | <sup>19</sup> |
| Colorectal Cancer | Colorectal | 121 | 73 | 48 | <sup>20</sup> |

Two types of microbiome profiles were extracted from the metagenomic samples: 1) strain-level marker profile and 2) species-level relative abundance profile. MetaPhlAn2 was utilized to extract these profiles with default parameters ^7^. We utilized MetAML to preprocess the abundance profile by selecting species-level features and excluding sub-species-level features ^12^. The strain-level marker profile consists of binary values indicating the presence (1) or absence (0) of a certain strain. The species-level relative abundance profile consists of real values in [0,1] indicating the percentages of the species in the total observed species. The abundance profile has a few hundred dimensions, whereas the marker profile has a much larger number of dimensions, up to over a hundred thousand in the current data (Table 2).

**Table 2.** The number of dimensions of the preprocessed microbiome profiles

| Profile type | IBD | EW-T2D | C-T2D | Obesity | Cirrhosis | Colorectal |
| --- | --- | --- | --- | --- | --- | --- |
| marker profile | 91,756 | 83,456 | 119,792 | 99,568 | 120,553 | 108,034 |
| abundance profile | 443 | 381 | 572 | 465 | 542 | 503 |

### Deep Representation Learning

An autoencoder is a neural network reconstructing its input *x*. Internally, its general form consists of an encoder function *f*_*ϕ*_(·) and a decoder function *f*’_*θ*_(·) where *ϕ* and *θ* are parameters of encoder and decoder functions, respectively. An autoencoder is trained to minimize the difference between an input *x* and a reconstructed input *x*’, the reconstruction loss (e.g., squared error) that can be written as follows:

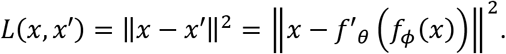

After training an autoencoder, we are interested in obtaining a latent representation *z* = *f*_*ϕ*_(*x*) of the input using the trained encoder. The latent representation, usually in a much lower-dimensional space than the original input, contains sufficient information for reconstructing the original input as close as possible. We utilized this representation to train classifiers for disease prediction.

For the DeepMicro framework, we incorporated various deep representation learning techniques, including shallow autoencoder (SAE), deep autoencoder (DAE), variational autoencoder (VAE), and convolutional autoencoder (CAE), to learn a low-dimensional embedding for microbiome profiles. Note that the diverse combinations of hyper-parameters defining the structure of autoencoders (e.g., the number of units and layers) have been explored in a grid fashion as described below, however, users are not limited to the tested hyper-parameters and can use their own hyper-parameter grid fitted to their data.

Firstly, we utilized SAE, the simplest autoencoder structure composed of the encoder part where the input layer is fully connected with the latent layer, and the decoder part where the output layer produces reconstructed input *x*’ by taking weighted sums of outputs of the latent layer. We introduced a linear activation function for the latent and output layer. Other options for the loss and activation functions are available for users (such as binary cross-entropy and sigmoid function). Initial values of the weights and bias were initialized with Glorot uniform initializer ^21^. We examined five different sizes of dimensions for the latent representation (32, 64, 128, 256, and 512).

In addition to the SAE model, we implemented the DAE model by introducing hidden layers between the input and latent layers as well as between the latent and output layers. All of the additional hidden layers were equipped with Rectified Linear Unit (ReLu) activation function and Glorot uniform initializer. The same number of hidden layers (one layer or two layers) were inserted into both encoder and decoder parts. Also, we gradually increased the number of hidden units. The number of hidden units in the added layers was set to the double of the successive layer in the encoder part and to the double of the preceding layer in the decoder part. With this setting, model complexity is controlled by both the number of hidden units and the number of hidden layers, maintaining structural symmetry of the model. For example, if the latent layer has 512 hidden units and if two layers are inserted to the encoder and decoder parts, then the resulting autoencoder has 5 hidden layers with 2048, 1024, 512, 1024, and 2048 hidden units, respectively. Similar to SAE, we varied the number of hidden units in the latent layer as follows: 32, 64, 128, 256, 512, thus, in total, we tested 10 different DAE architectures (Table S2).

A variational autoencoder (VAE) learns probabilistic representations *z* given input *x* and then use these representations to reconstruct input *x*’ ^22^. Using variational inference, the true posterior distribution of latent embeddings (i.e., *p*(*z*|*x*)) can be approximated by the introduced posterior *q*_*ϕ*_(*z*|*x*) where *ϕ* are parameters of an encoder network. Unlike the previous autoencoders learning an unconstrained representation, VAE learns a generalized latent representation under the assumption that the posterior approximation follows Gaussian distribution. The encoder network encodes the means and variances of the multivariate Gaussian distribution. The latent representation *z* can be sampled from the learned posterior distribution *q*_*ϕ*_(*z*|*x*) ~ N(*μ*, Σ). Then the sampled latent representation is passed into the decoder network to generate the reconstructed input *x*’ ~ *g*_*θ*_(*x*|*z*) where *θ* are the parameters of the decoder.

To approximate the true posterior, we need to minimize the Kullback-Leibler (KL) divergence between the introduced posterior and the true posterior,

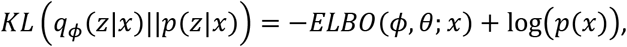

rewritten as

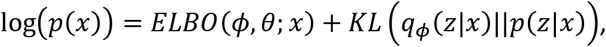

where *ELBO*(*ϕ,θ,x*) is an evidence lower bound on the log probability of the data because the KL term must be greater than or equal to zero. It is intractable to compute the KL term directly but minimizing the KL divergence is equivalent to maximizing the lower bound, decomposed as follows:

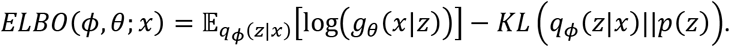

The final objective function can be induced by converting the maximization problem to the minimization problem.

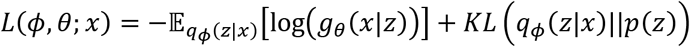

The first term can be viewed as a reconstruction term as it forces the inferred latent representation to recover its corresponding input and the second KL term can be considered as a regularization term to modulate the posterior of the learned representation to be Gaussian distribution. We used ReLu activation and Glorot uniform initializer for intermediate hidden layers in encoder and decoder. One intermediate hidden layer was used and the number of hidden units in it varied from 32, 64, 128, 256, to 512. The latent layer was set to 4, 8, or 16 units. Thus, altogether we tested 15 different model structures.

Instead of fully connected layers, a convolutional autoencoder (CAE) is equipped with convolutional layers in which each unit is connected to only local regions of the previous layer ^23^. A convolutional layer consists of multiple filters (kernels) and each filter has a set of weights used to perform convolution operation that computes dot products between a filter and a local region ^24^. We used ReLu activation and Glorot uniform initializer for convolutional layers. We did not use any pooling layer as it may generalize too much to reconstruct an input. The n-dimensional input vector was reshaped like a squared image with a size of *d* × *d* × 1 where 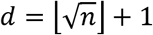. As *d*^2^ ≥ *n*, we padded the rest part of the reshaped input with zeros. To be flexible to an input size, the filter size of the first convolutional layer was set to 10% of the input width and height, respectively (i.e. [0.1*d*] × [0.1*d*]). For the first convolutional layer, we used 25% of the filter size as the size of stride which configures how much we slide the filter. For the following convolutional layers in the encoder part, we used 10% of the output size of the preceding layer as the filter size and 50% of this filter size as the stride size. All units in the last convolutional layer of the encoder part have been flattened in the following flatten layer which is designated as a latent layer. We utilized convolutional transpose layers (deconvolutional layers) to make the decoder symmetry to the encoder. In our experiment, the number of filters in a convolutional layer was set to half of that of the preceding layer for the encoder part. For example, if the first convolutional layer has 64 filters and there are three convolutional layers in the encoder, then the following two convolutional layers have 32 and 16 filters, respectively. We varied the number of convolutional layers from 2 to 3 and tried five different numbers of filters in the first convolutional layer (4, 8, 16, 32, and 64). In total, we tested 10 different CAE model structures.

To train deep representation models, we split each dataset into a training set, a validation set, and a test set (64% training set, 16% validation set, and 20% test set; Figure S1). Note that the test set was withheld from training the model. We used the early-stopping strategy, that is, trained the models on the training set, computed the reconstruction loss for the validation set after each epoch, stopped the training if there was no improvement in validation loss during 20 epochs, and then selected the model with the least validation loss as the best model. We used mean squared error for reconstruction loss and applied adaptive moment estimation (Adam) optimizer for gradient descent with default parameters (learning rate: 0.001, epsilon: 1e-07) as provided in the original paper ^25^. We utilized the encoder part of the best model to produce a low-dimensional representation of the microbiome data for downstream disease prediction.

### Prediction of disease states based on the learned representation

We built classification models based on the encoded low-dimensional representations of microbiome profiles (Figure 1). Three machine learning algorithms, support vector machine (SVM), random forest (RF), and Multi-Layer Perceptron (MLP), were used. We explored hyper-parameter space with grid search. SVM maximizes the margin between the supporting hyperplanes to optimize a decision boundary separating data points of different classes ^26^. In this study, we utilized both radial basis function (RBF) kernel and a linear kernel function to compute decision margins in the transformed space to which the original data was mapped. We varied penalty parameter *C* (2^−5^, 2^−3^, …, 2^5^) for both kernels as well as kernel coefficient *gamma* (2^−15^, 2^−13^, …, 2^3^) for RBF kernel. In total, 60 different combinations of hyper-parameters were examined to optimize SVM (Table S2).

**Figure 1.**
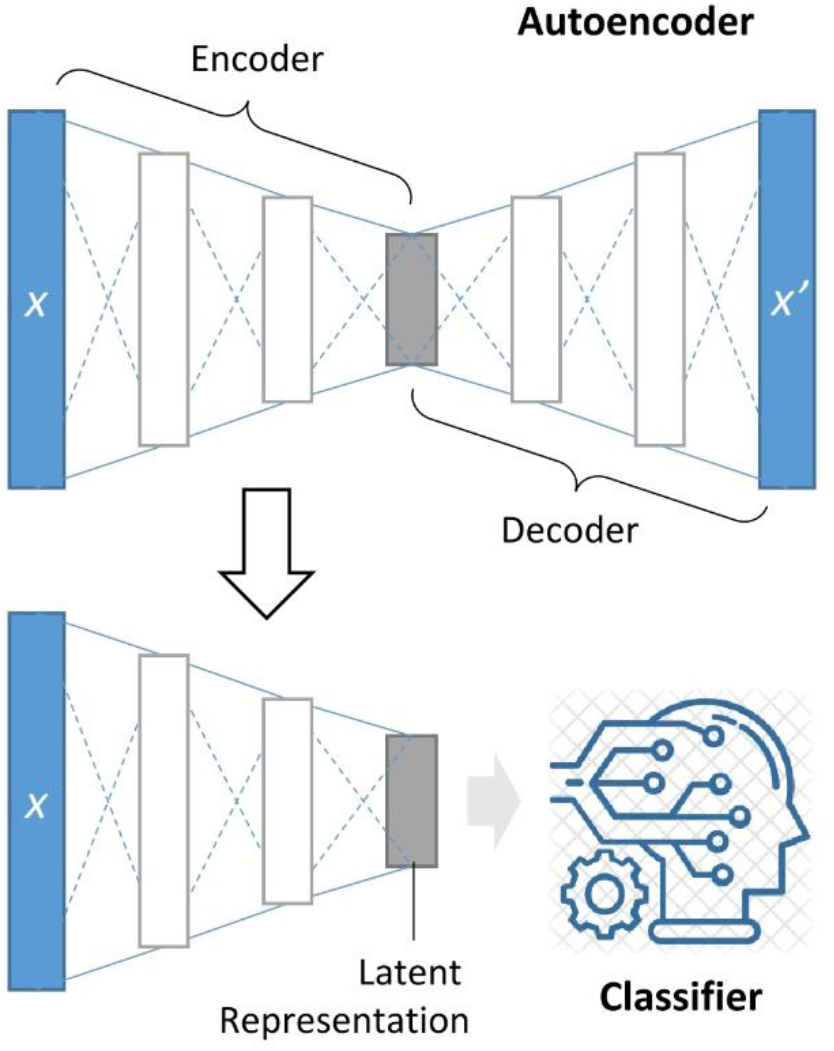
DeepMicro framework. An autoencoder is trained to map the input X to the low-dimensional latent space with the encoder and to reconstruct X with the decoder. The encoder part is reused to produce a latent representation of any new input X that is in turn fed into a classification algorithm to determine whether the input is the positive or negative class.

RF builds multiple decision trees based on various sub-samples of the training data and merges them to improve the prediction accuracy. The size of sub-samples is the same as that of training data but the samples are drawn randomly with replacement from the training data. For the hyper-parameter grid of RF classifier, the number of trees (estimators) was set to 100, 300, 500, 700, and 900, and the minimum number of samples in a leaf node was altered from 1 to 5. Also, we tested two criteria, Gini impurity and information gain, for selecting features to split a node in a decision tree. For the maximum number of features considered to find the best split at each split, we used a square root of n and a logarithm to base 2 of *n* (*n* is the sample size). In total, we tested 100 combinations of hyper-parameters of RF.

MLP is an artificial neural network classifier that consists of an input layer, hidden layers, and an output layer. All of the layers are fully connected to their successive layer. We used ReLu activations for all hidden layers and sigmoid activation for the output layer that has a single unit. The number of units in the hidden layers was set to half of that of the preceding layer except the first hidden layer. We varied the number of hidden layers (1, 2, and 3), the number of epochs (30, 50, 100, 200, and 300), the number of units in the first hidden layer (10, 30, 50, 100), and dropout rate (0.1 and 0.3). In total, 120 hyper-parameter combinations were tested in our experiment.

We implemented DeepMicro in Python 3.5.2 using machine learning and data analytics libraries, including Numpy 1.16.2, Pandas 0.24.2, Scipy 1.2.1, Scikt-learn 0.20.3, Keras 2.2.4, and Tensorflow 1.13.1. Source code is publicly available at the git repository (https://github.com/minoh0201/DeepMicro).

### Performance Evaluation

To avoid an overestimation of prediction performance, we designed a thorough performance evaluation scheme (Figure S1). For a given dataset (e.g. Cirrhosis), we split it into training and test set in the ratio of 8:2 with a given random partition seed, keeping a ratio between classes in both training and test set to be the same as that of the given dataset. Using only the training set, a representation learning model was trained. Then, the learned representation model was applied to the training set and test set to obtain dimensionality-reduced training and test set. After the dimensionality has been reduced, we conducted 5-fold cross-validation on the training set by varying hyper-parameters of classifiers. The best hyper-parameter combination for each classifier was selected by averaging an accuracy metric of the five different results. The area under the receiver operating characteristics curve (AUC) was used for performance evaluation. We trained a final classification model using the whole training set with the best combination of hyper-parameters and tested it on the test set. This procedure was repeated five times by changing the random partition seed at the beginning of the procedure. The resulting AUC scores were averaged and the average was used to compare model performance.

## Results

We developed DeepMicro, a deep representation learning framework for predicting individual phenotype based on microbiome profiles. Various autoencoders (SAE, DAE, VAE, and CAE) have been utilized to learn a low-dimensional representation of the microbiome profiles. Then three classification models including SVM, RF, and MLP were trained on the learned representation to discriminate between disease and control sample groups. We tested our framework on six disease datasets (Table 1), including inflammatory bowel disease (IBD), type 2 diabetes in European women (EW-T2D), type 2 diabetes in Chinese (C-T2D), obesity (Obesity), liver cirrhosis (Cirrhosis), and colorectal cancer (Colorectal). For all the datasets, two types of microbiome profiles, strain-level marker profile and species-level relative abundance profile, have been extracted and tested (Table 2). Also, we devised a thorough performance evaluation scheme that isolates the test set from the training and validation sets in the hyper-parameter optimization phase to compare various models (See Methods and Figure S1).

We compared our method to the current best approach (MetAML) that directly trained classifiers, such as SVM and RF, on the original microbiome profile ^12^. We utilized the same hyper-parameters grid used in MetAML for each classification algorithm. In addition, we tested Principal Component Analysis (PCA) and Gaussian Random Projection (RP), using them as the replacement of the representation learning to observe how traditional dimensionality reduction algorithms behave. For PCA, we selected the principal components explaining 99% of the variance in the data ^27^. For RP, we set the number of components to be automatically adjusted according to Johnson-Lindenstrauss lemma (eps parameter was set to 0.5) ^28–30^.

We picked the best model for each approach in terms of prediction performance and compared the approaches across the datasets. Figure 2 shows the results of DeepMicro and the other approaches for the strain-level marker profile. DeepMicro outperforms the other approaches for five datasets, including IBD (AUC = 0.955), EW-T2D (AUC = 0.899), C-T2D (AUC = 0.763), Obesity (AUC = 0.659), and Cirrhosis (AUC = 0.940). For Colorectal dataset, DeepMicro has slightly lower performance than the best approach (DeepMicro’s AUC = 0.803 vs. MetAML’s AUC = 0.811). The marker profile-based models generally perform better than the abundance profile-based models (Figure S8 and S2). The only exception is Obesity dataset for which the abundance-based DeepMicro model shows better performance (AUC = 0.674). Note that as AUC could be misleading in an imbalanced classification scenario ^31^, we also evaluated the area under the precision-recall curve (AUPRC) for the imbalanced data set IBD and observed the same trend between AUC and AUPRC (Table S3).

**Figure 2.**
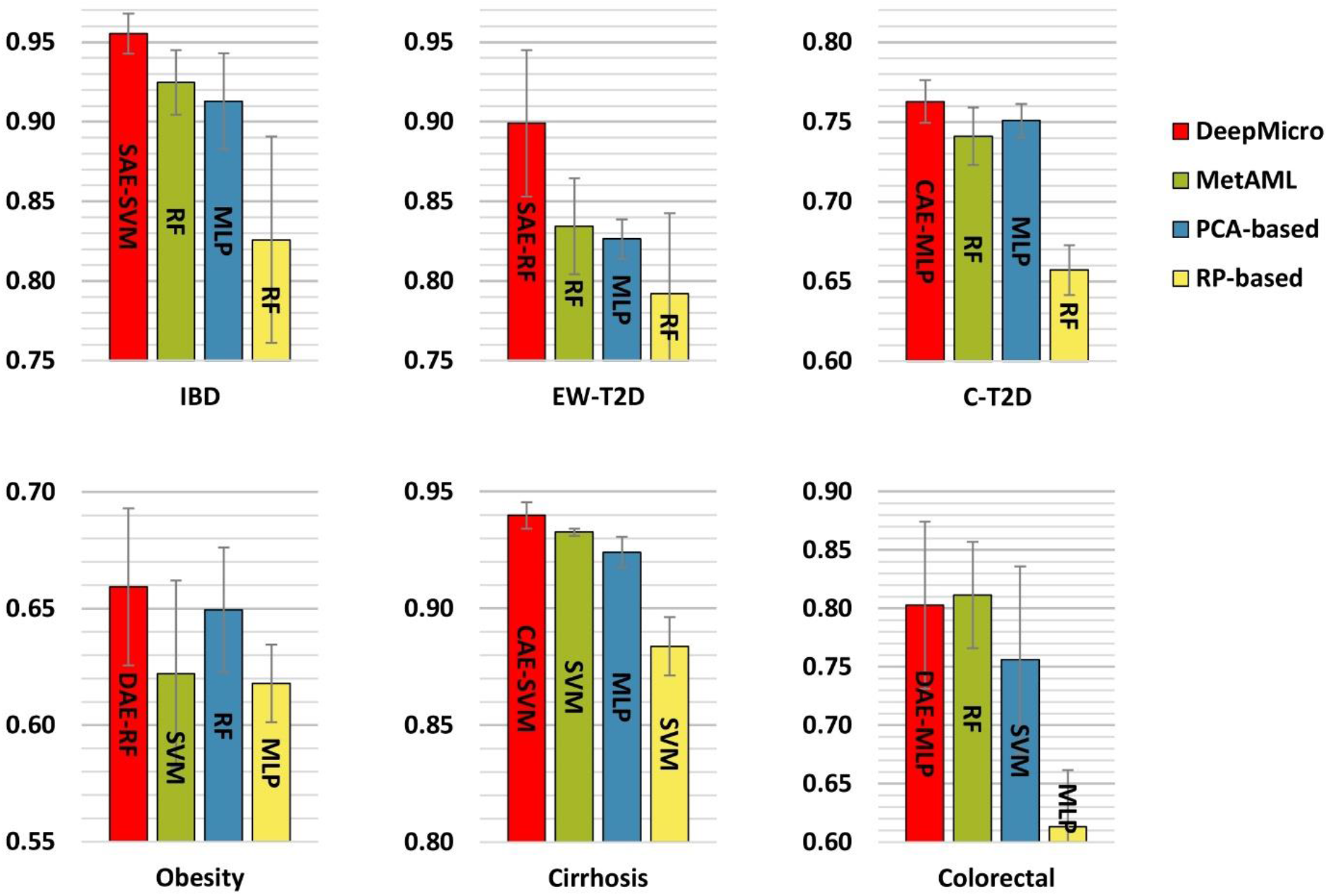
Disease prediction performance for marker profile-based models. Prediction performance of various methods built on marker profile has been assessed with AUC. MetAML utilizes support vector machine (SVM) and random forest (RF), and the superior model is presented (green). Principal component analysis (PCA; blue) and gaussian random projection (RP; yellow) have been applied to reduce dimensions of datasets before classification. DeepMicro (red) applies shallow autoencoder (SAE), deep autoencoder (DAE), variational autoencoder (VAE), and convolutional autoencoder (CAE) for dimensionality reduction. Then SVM, RF, and multi-layer perceptron (MLP) classification algorithms have been used.

For marker profile, none of the autoencoders dominate across the datasets in terms of getting the best representation for classification. Also, the best classification algorithm varied according to the learned representation and to the dataset (Figure 3). For abundance profile, CAE dominates over the other autoencoders with RF classifier across all the datasets (Figure S3).

**Figure 3.**
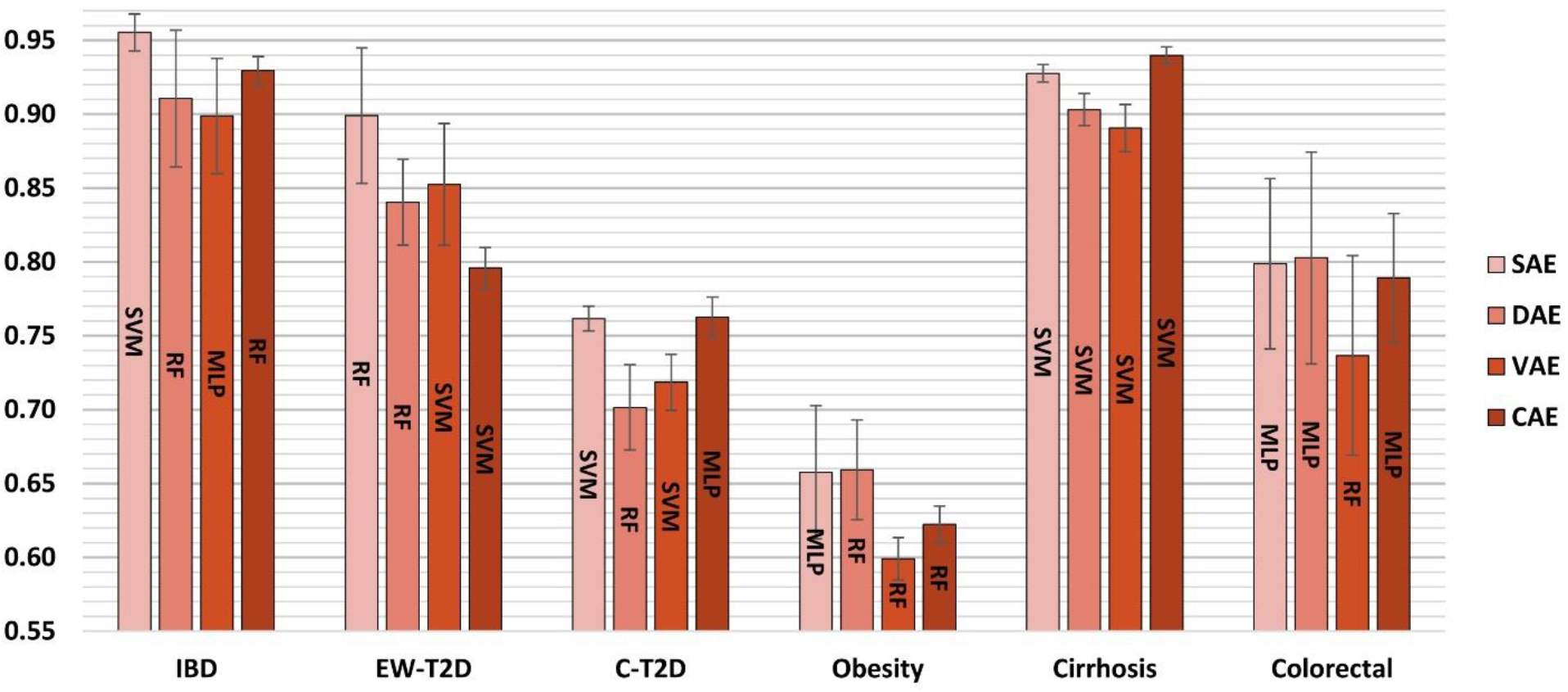
Disease prediction performance for different autoencoders based on marker profile (assessed with AUC). Classifiers used: support vector machine (SVM), random forest (RF), and multi-layer perceptron (MLP); Autoencoders used: shallow autoencoder (SAE), deep autoencoder (DAE), variational autoencoder (VAE), and convolutional autoencoder (CAE)

We also directly trained MLP on the dataset without representation learning and compared the prediction performance with that of the traditional approach (the best between SVM and RF). It is shown that MLP performs better than MetAML in three datasets, EW-T2D, C-T2D, and Obesity, when marker profile is used (Figure S4). However, when abundance profile is used, the performance of MLP was worse than that of the traditional approach across all the datasets (Figures S5).

Furthermore, we compared running time of DeepMicro on marker profiles with a basic approach not using representation learning. For comparison, we tracked both training time and representation learning time. For each dataset, we tested the best performing representation learning model producing the highest AUC score (i.e. SAE for IBD and EW-T2D, DAE for Obesity and Colorectal, and CAE for C-T2D and Cirrhosis; Table S1). We fixed the seed for random partitioning of the data, and applied the formerly used performance evaluation procedure where 5-fold cross-validation is conducted on the training set to obtain the best hyper-parameter with which the best model is trained on the whole training set and is evaluated on the test set (See Methods). The computing machine we used for timestamping is running on Ubuntu 18.04 and equipped with an Intel Core i9-9820X CPU (10 cores), 64 GB Memory, and a GPU of NVIDIA GTX 1080 Ti. We note that our implementation utilizes GPU when it learns representations and switches to CPU mode to exhaustively use multiple cores in a parallel way to find best hyper-parameters of the classifiers. Table 3 shows the benchmarking result on marker profile. It is worth noting that DeepMicro is 8X to 30X times faster than the basic approach (17X times faster on average). Even if MLP is excluded from the benchmarking because it requires heavy computation, DeepMicro is up to 5X times faster than the basic (2X times faster on average).

**Table 3.** Time benchmark for DeepMicro and basic approaches without representation learning (in sec)

| Method |  | IBD | EW-T2D | C-T2D | Obesity | Cirrhosis | Colorectal |
| --- | --- | --- | --- | --- | --- | --- | --- |
| Basic approach | **SVM\*** | 126 | 85 | 1705 | 711 | 777 | 187 |
|  | **RF** | 42 | 41 | 99 | 79 | 72 | 50 |
|  | **MLP** | 3,776 | 2,449 | 12,057 | 8,186 | 8,593 | 4,508 |
| **Total elapsed** |  | 3,943 | 2,575 | 13,861 | 8,976 | 9,442 | 4,745 |
| DeepMicro | **RL** | 74 | 194 | 554 | 113 | 521 | 215 |
|  | **SVM** | 2 | 2 | 8 | 8 | 17 | 2 |
|  | **RF** | 28 | 28 | 47 | 33 | 40 | 30 |
|  | **MLP** | 103 | 93 | 188 | 137 | 287 | 105 |
| **Total elapsed** |  | 207 | 317 | 798 | 291 | 864 | 352 |

\*RL: Representation Learning; SVM: Support Vector Machine; RF: Random Forest; MLP: Multi-layer Perceptron

## Discussion

We developed a deep learning framework transforming a high-dimensional microbiome profile into a low-dimensional representation and building classification models based on the learned representation. At the beginning of this study, the main goal was to reduce dimensions as strain-level marker profile has too many dimensions to handle, expecting that noisy and unnecessary information fades out and the refined representation becomes tractable for downstream prediction. Firstly, we tested PCA on marker profile and it showed a slight improvement in prediction performance for C-T2D and Obesity but not for the others. The preliminary result indicates that either some of the meaningful information was dropped or noisy information still remains. To learn meaningful feature representations, we trained various autoencoders on microbiome profiles. Our intuition behind the autoencoders was that the learned representation should keep essential information in a condensed way because autoencoders are forced to prioritize which properties of the input should be encoded during the learning process. We found that although the most appropriate autoencoder usually allows for better representation that in turn results in better prediction performance, what kind of autoencoder is appropriate highly depends on problem complexity and intrinsic properties of the data.

In the previous study, it has been shown that adding healthy controls of the other datasets could improve prediction performance assessed by AUC ^12^. To check if this finding can be reproduced, for each dataset, we added control samples of the other datasets only into the training set and kept the test set the same as before. Figure S6 shows the difference between the best performing models built with and without additional controls. In general, prediction performance dropped (on average by 0.037) once negative (control) samples are introduced to the training set across the datasets in almost all approaches except only a few cases (Figure S6). In contrast to the previous study, the result indicates that the insertion of only negative samples into the training set may not help to improve the classification models, and a possible explanation might be that changes in the models rarely contribute to improving the classification of positive samples ^32^. Interestingly, if we added negative samples into the whole data set before split it into training and test set, we usually observed improvements in prediction performance. However, we found that these improvements are trivial because introducing negative samples into the test set easily reduces false positive rate (as the denominator of false positive rate formula is increased), resulting in higher AUC scores.

Even though adding negative samples might not be helpful for a better model, it does not mean that additional samples are meaningless. We argue that more samples can improve prediction performance, especially when a well-balanced set of samples is augmented. To test this argument, we gradually increased the proportion of the training set and observed how prediction performance changed over the training sets of different sizes. Generally, improved prediction performance has been observed as more data of both positive and negative samples are included (Figure S7). With the continued availability of large samples of microbiome data, the deep representation learning framework is expected to become increasingly effective for both condensed representation of the original data and also downstream prediction based on the deep representation.

DeepMicro is publicly available software which offers cutting-edge deep learning techniques for learning meaningful representations from the given data. Researchers can apply DeepMicro to their high-dimensional microbiome data to obtain a robust low-dimensional representation for the subsequent supervised or unsupervised learning. For predictive problems increasingly studied with microbiome data such as drug response prediction, forensic human identification, and food allergy prediction, deep representation learning might be useful in terms of boosting the model performance. Moreover, it might be worthwhile to use the learned representation for clustering analysis. The distance between data points in the latent space can be a basis for clustering microbiome samples and it could help capture the shared characteristics within a group which are difficult to be identified in the original data space. DeepMicro has been used to deal with microbiome data but it is not limited to a specific type of data and its application can be extended to various omics data, such as genome and proteome data.

## Data availability

All data and codes are available at https://github.com/minoh0201/DeepMicro.

## List of abbreviations

IBD: inflammatory bowel disease
EW-T2D: type 2 diabetes in European women
C-T2D: type 2 diabetes in Chinese
Obesity: obesity
Cirrhosis: liver cirrhosis
Colorectal: colorectal cancer
SAE: shallow autoencoder
DAE: deep autoencoder
VAE: variational autoencoder
CAE: convolutional autoencoder
ReLu: rectified linear unit
KL: Kullback-Leibler
SVM: support vector machine
RF: random forest
MLP: multi-layer perceptron
RBF: radial basis function
AUC: area under the receiver operating characteristics curve
PCA: Principal Component Analysis
RP: Gaussian Random Projection

## Acknowledgments

This work is partially supported by the funding from Data and Decisions Destination Area at Virginia Tech. Also, we thank Dr. Bert Huang for the constructive discussion.

## Author contributions

MO designed the study, collected data, implemented the software, and performed experiments. MO and LZ interpreted the results and wrote the manuscript. All authors read and approved the final manuscript.

