## Supplementary material for "DeepMicro: deep representation learning for disease prediction based on microbiome data": Figure S

Min Oh[1] and Liqing Zhang[1,*]

[1]Department of Computer Science, Virginia Tech, Blacksburg, VA, USA

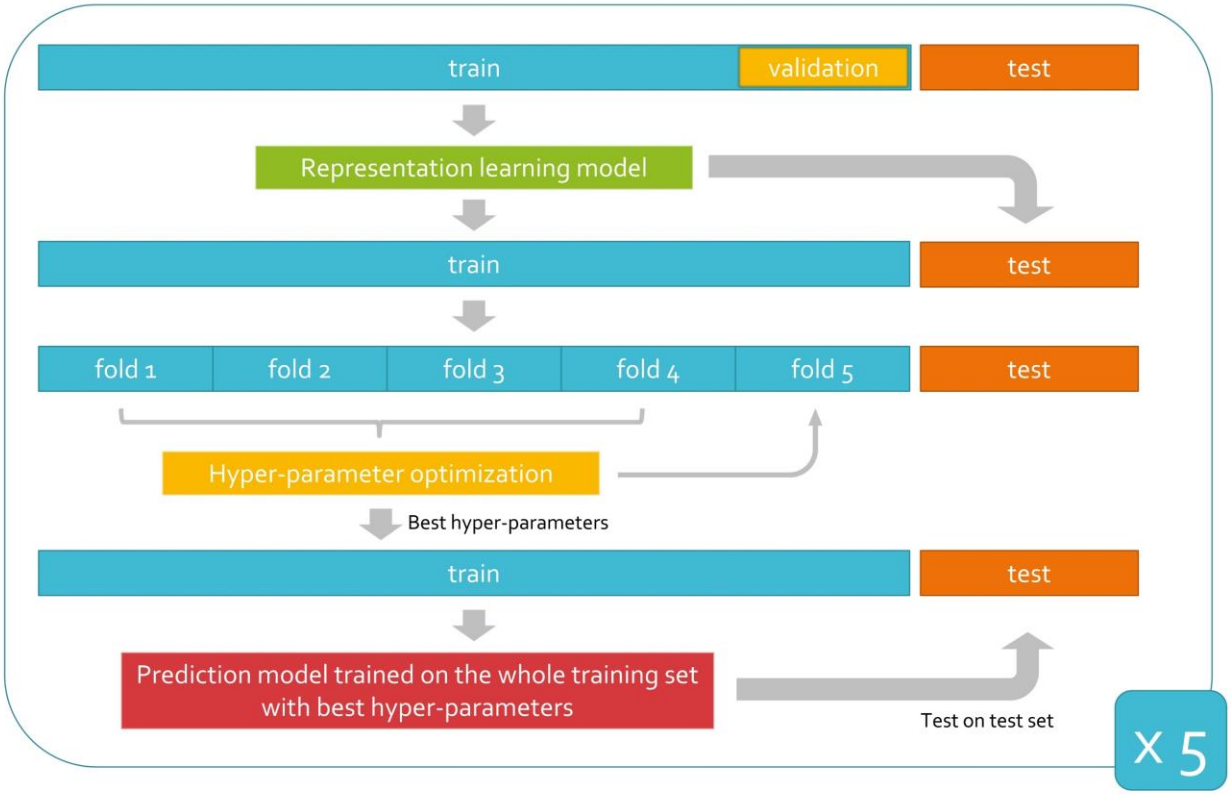


**Figure S1.** Performance evaluation scheme

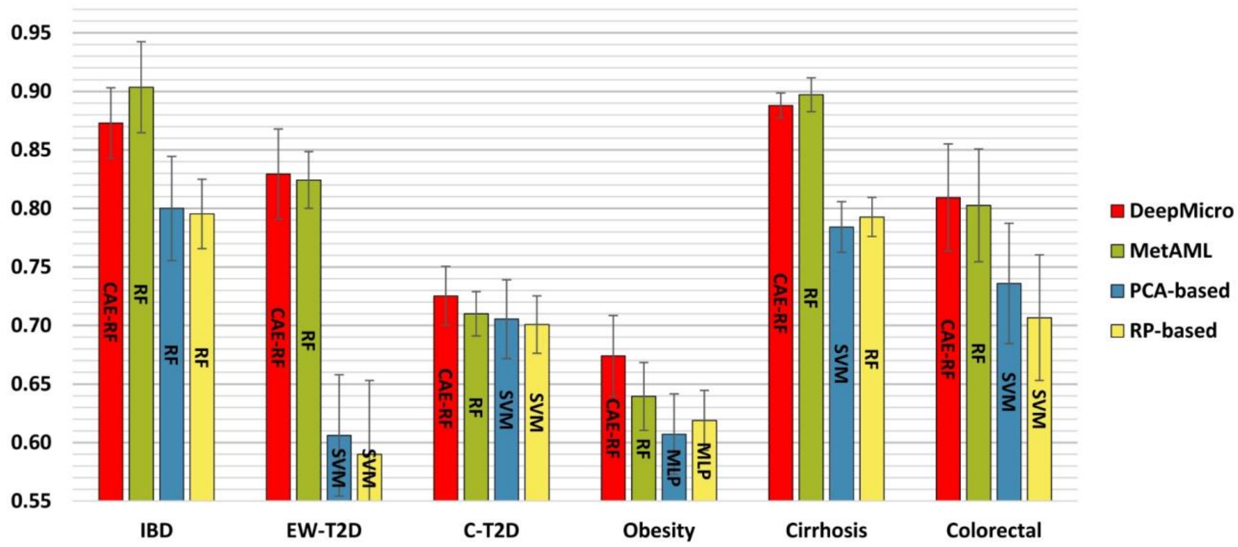


**Figure S2.** Disease prediction performance for abundance profile-based models. Prediction performance of various methods built on marker profile has been assessed with AUC. MetAML utilizes support vector machine (SVM) and random forest (RF), and the superior model is presented (green). Principal component analysis (PCA; blue) and gaussian random projection (RP; yellow) have been applied to reduce dimensions of datasets before classification. DeepMicro (red) applies shallow autoencoder (SAE), deep autoencoder (DAE), variational autoencoder (VAE), and convolutional autoencoder (CAE) for dimensionality reduction. Then SVM, RF, and multi-layer perceptron (MLP) classification algorithms have been used.

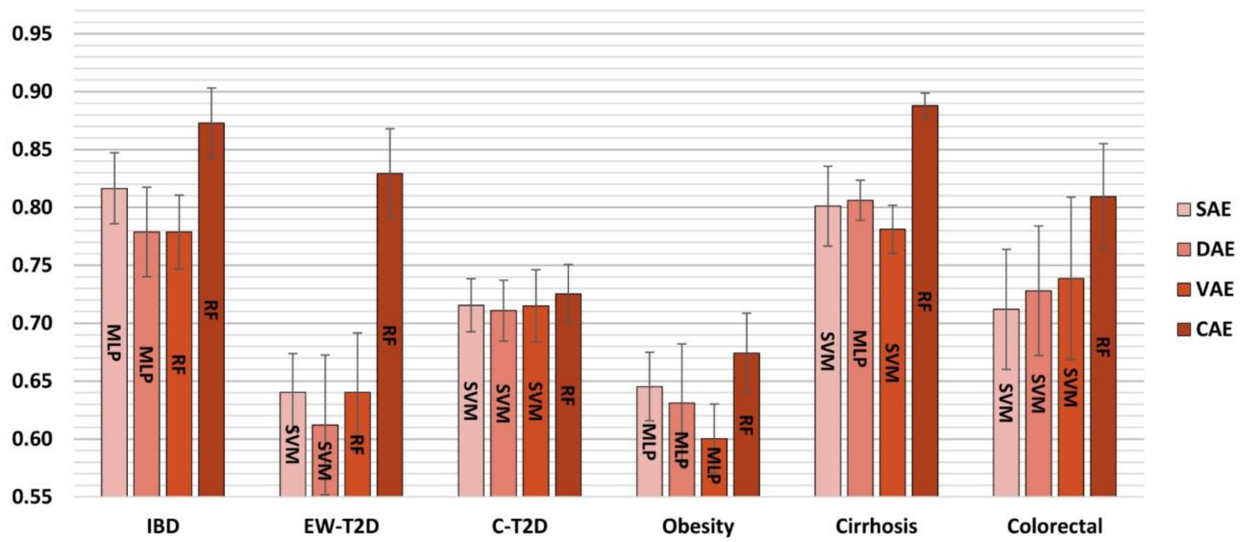


**Figure S3.** Disease prediction performance for different autoencoders based on abundance profile (assessed with AUC). Classifiers used: support vector machine (SVM), random forest (RF), and multi-layer perceptron (MLP); Autoencoders used: shallow autoencoder (SAE), deep autoencoder (DAE), variational autoencoder (VAE), and convolutional autoencoder (CAE)

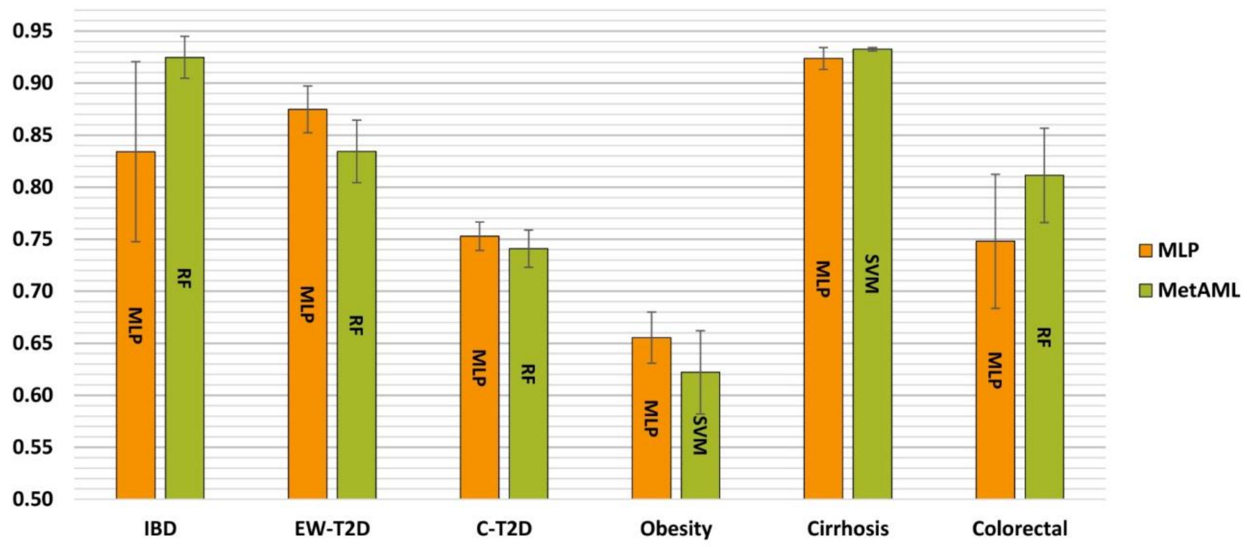


**Figure S4.** Disease prediction performance of multi-layer perceptron without representation learning based on marker profile

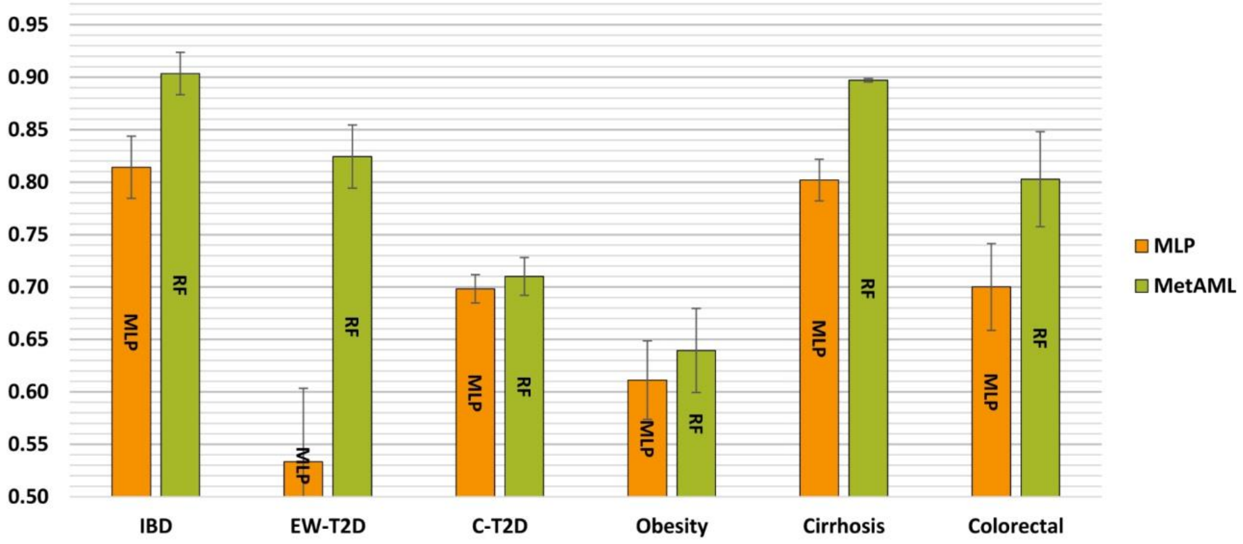


**Figure S5.** Disease prediction performance of multi-layer perceptron without representation learning based on abundance profile

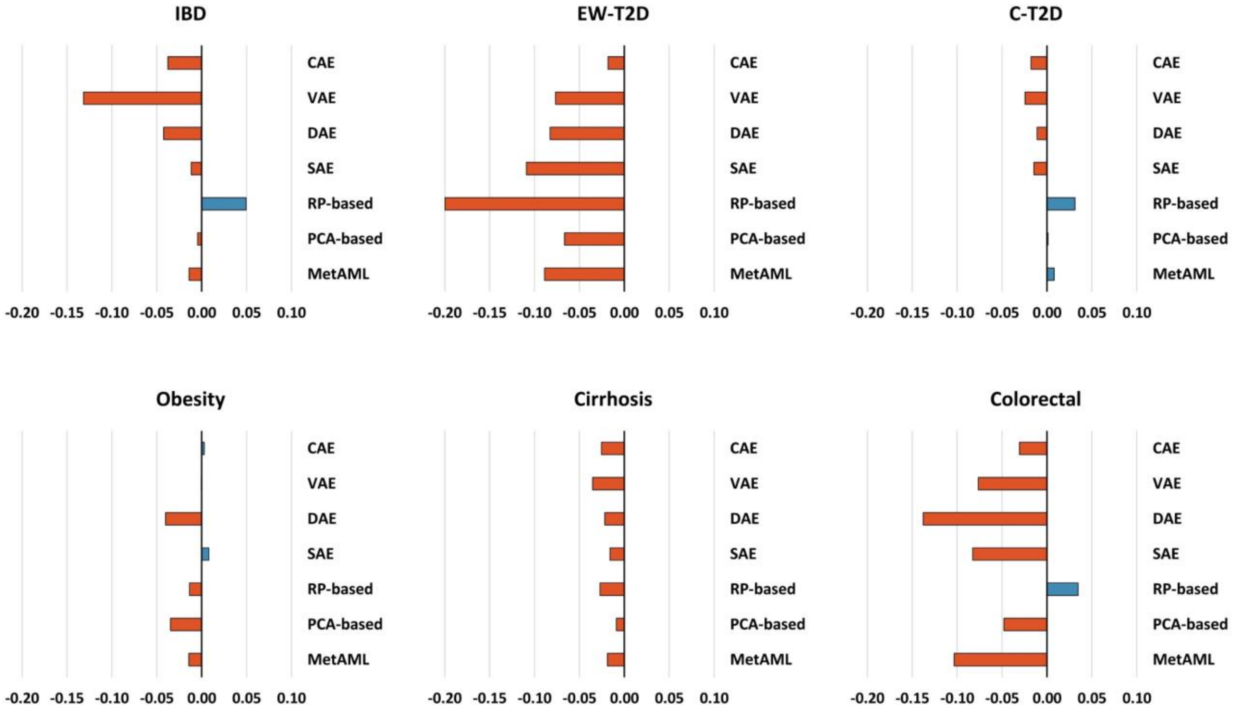


**Figure S6.** Impact of introducing negative samples into the training set on AUC

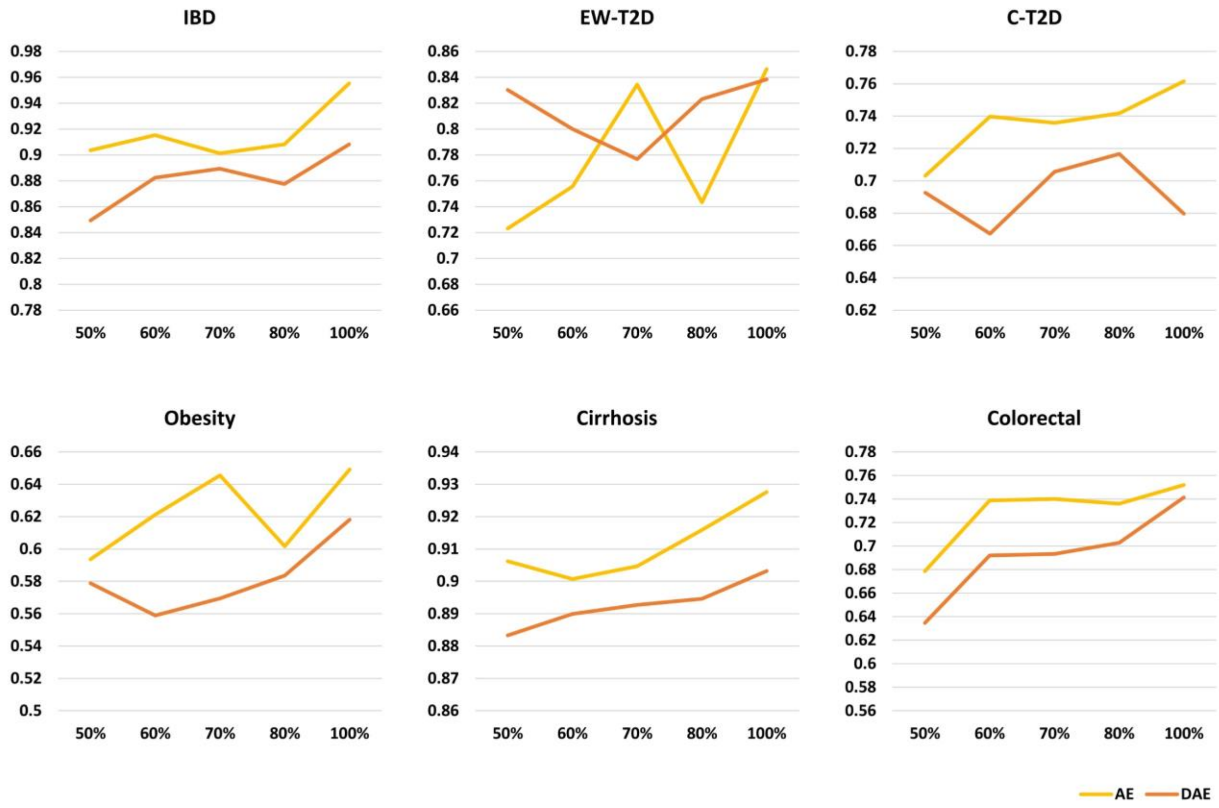


**Figure S7.** Prediction performance changes over the increasing data points in the training set

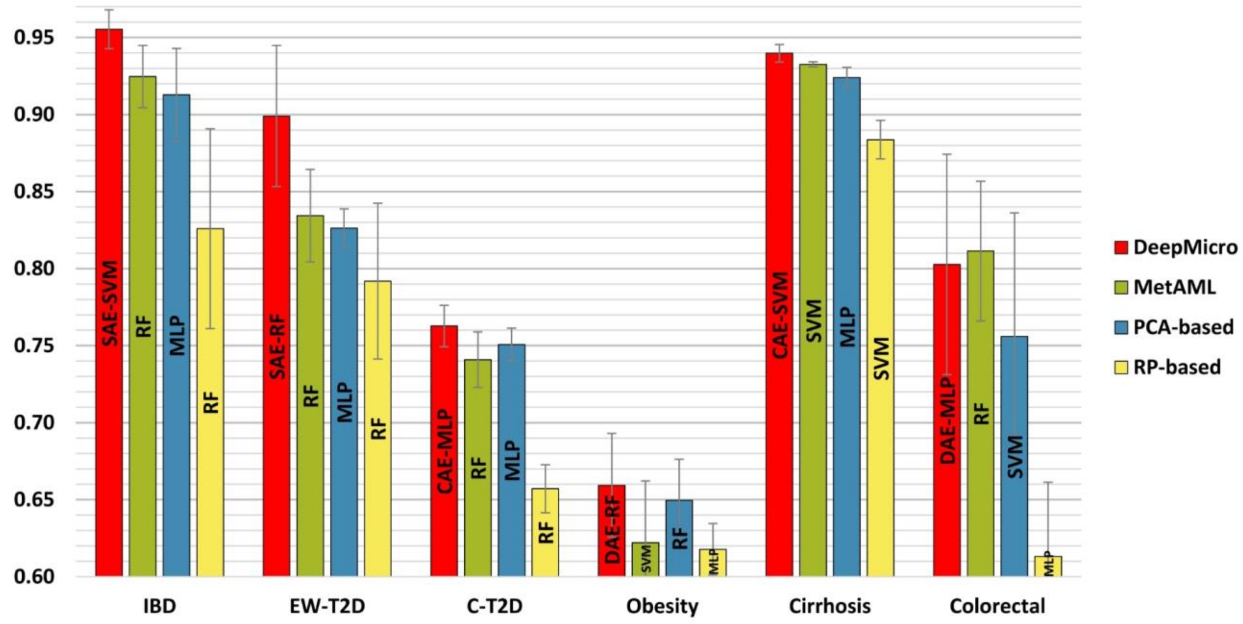


**Figure S8.** Disease prediction performance for marker profile-based models (fixed scale).

**Table S1.** The best representation learning model structures for each dataset

| Microbiome Profile Type | Dataset | Size of Original Dim[#] | Representation Learning Model | Encoder Structure* | Size of Latent Dim | Classifier | Averaged AUC (Standard Error) |
|---|---|---|---|---|---|---|---|
| Strain-level marker profile | IBD | 91,756 | SAE | 64 | 64 | SVM | **0.955 (0.013)** |
| | | | DAE | 512-256-128 | 128 | RF | 0.911 (0.046) |
| | | | VAE | 128-4 | 4 | MLP | 0.899 (0.039) |
| | | | CAE | 8-4 | 1,936 | RF | 0.929 (0.010) |
| | EW-T2D | 83,456 | SAE | 256 | 256 | RF | **0.899 (0.046)** |
| | | | DAE | 256-128-64 | 64 | RF | 0.840 (0.029) |
| | | | VAE | 256-16 | 16 | SVM | 0.853 (0.041) |
| | | | CAE | 8-4 | 1,764 | SVM | 0.796 (0.014) |
| | C-T2D | 119,792 | SAE | 512 | 512 | SVM | 0.762 (0.008) |
| | | | DAE | 256-128 | 128 | RF | 0.702 (0.029) |
| | | | VAE | 128-16 | 16 | SVM | 0.719 (0.019) |
| | | | CAE | 4-2 | 968 | MLP | **0.763 (0.014)** |
| | Obesity | 99,568 | SAE | 512 | 512 | MLP | 0.658 (0.045) |
| | | | DAE | 256-128 | 128 | RF | **0.659 (0.034)** |
| | | | VAE | 512-8 | 8 | RF | 0.599 (0.014) |
| | | | CAE | 64-32 | 16,928 | RF | 0.622 (0.012) |
| | Cirrhosis | 120,553 | SAE | 256 | 256 | SVM | 0.928 (0.006) |
| | | | DAE | 512-256-128 | 128 | SVM | 0.903 (0.011) |
| | | | VAE | 256-8 | 8 | SVM | 0.891 (0.016) |
| | | | CAE | 16-8 | 3,872 | SVM | **0.940 (0.006)** |
| | Colorectal | 108,034 | SAE | 32 | 32 | MLP | 0.799 (0.058) |
| | | | DAE | 512-256-128 | 128 | MLP | **0.803 (0.072)** |
| | | | VAE | 256-8 | 8 | RF | 0.737 (0.068) |
| | | | CAE | 4-2-1 | 441 | MLP | 0.789 (0.044) |
| Species-level relative abundance profile | IBD | 443 | SAE | 512 | 512 | MLP | 0.817 (0.031) |
| | | | DAE | 512-256 | 256 | MLP | 0.779 (0.039) |
| | | | VAE | 32-8 | 8 | RF | 0.779 (0.032) |
| | | | CAE | 32-16-8 | 3,872 | RF | **0.873 (0.030)** |
| | EW-T2D | 381 | SAE | 256 | 256 | SVM | 0.640 (0.033) |
| | | | DAE | 1024-512 | 512 | SVM | 0.612 (0.060) |
| | | | VAE | 64-8 | 8 | RF | 0.640 (0.051) |
| | | | CAE | 16-8 | 3,200 | RF | **0.829 (0.039)** |
| | C-T2D | 572 | SAE | 64 | 64 | SVM | 0.715 (0.023) |
| | | | DAE | 128-64 | 64 | SVM | 0.711 (0.026) |
| | | | VAE | 512-16 | 16 | SVM | 0.715 (0.031) |
| | | | CAE | 4-2-1 | 576 | RF | **0.725 (0.025)** |
| | Obesity | 465 | SAE | 128 | 128 | MLP | 0.645 (0.030) |
| | | | DAE | 1024-512 | 512 | MLP | 0.631 (0.051) |
| | | | VAE | 256-4 | 4 | MLP | 0.600 (0.030) |
| | | | CAE | 4-2 | 968 | RF | **0.674 (0.034)** |
| | Cirrhosis | 542 | SAE | 32 | 32 | SVM | 0.801 (0.035) |
| | | | DAE | 1024-512 | 512 | MLP | 0.806 (0.017) |
| | | | VAE | 512-8 | 8 | SVM | 0.781 (0.021) |
| | | | CAE | 16-8-4 | 1,461 | RF | **0.888 (0.011)** |
| | Colorectal | 503 | SAE | 256 | 256 | SVM | 0.712 (0.052) |
| | | | DAE | 256-128 | 128 | SVM | 0.728 (0.056) |
| | | | VAE | 512-8 | 8 | SVM | 0.739 (0.070) |
| | | | CAE | 8-4 | 2,116 | RF | **0.809 (0.046)** |

*The number of units for SAE, DAE, and VAE; The number of filters for CAE; Layers are separated by a delimiter "-"

[#]Dim: Dimension; SAE: Sallow Autoencoder; DAE: Deep Autoencoder; VAE: Variational autoencoder; CAE: Convolutional autoencoder; SVM: Support Vector Machine; RF: Random Forest; MLP: Multi-layer Perceptron
